# Semaglutide Activates the Orexin/Hypocretin and Basal Forebrain Cholinergic Systems and Increases Acetylcholine Levels in the Hippocampus of Young and Aged Rats

**DOI:** 10.64898/2026.08.03.742612

**Authors:** Christian Wohlfeld, Alexis Blas, Jennifer Woodruff, Marla Frick, Natalia Maciejewska, Ayesha Patel, Claudia Grillo, Lawrence Reagan, Jim Fadel

**Affiliations:** University of South Carolina School of Medicine, Department of Pharmacology, Physiology and Neuroscience; WJB Dorn Veterans Affairs Medical Center

## Abstract

GLP-1 agonist drug repurposing efforts may establish new clinical niches in managing psychiatric and neurological disorders. However, a comprehensive understanding of GLP-1 neurobiology and an appreciation of specific neural mechanisms by which GLP-1 agonists might provide therapeutic effects is limited and stands as a barrier to these efforts. When considering current preclinical research evaluating GLP-1 agonist central mechanisms, a considerable knowledge gap remains regarding which specific cellular populations and systems define potential therapeutic effects in the brain. In this research, we used cFos immunohistochemistry to identify specific neuronal populations that exhibited altered cellular activity following acute administration of semaglutide to rats. We found that the orexin/hypocretin and basal forebrain cholinergic systems are activated following acute semaglutide administration in male and female young adult rats (3-5 months). Informed by the results of our histological analysis, we next employed in vivo microdialysis to test our hypothesis that semaglutide would acutely increase acetylcholine release in the rodent hippocampus. Here, we report that semaglutide acutely increases acetylcholine efflux in the ventral hippocampus of conscious and freely moving rats regardless of biological sex in both young adult and aged rats (23-26 months). Given the relevance of hippocampal cholinergic neurotransmission in learning and memory, our research mechanistically connects GLP-1 agonists with established targets in cognitive decline and dementia.

## Introduction

Glucagon-like peptide-1 receptor agonists (GLP-1 agonists) have proven to be highly clinically useful in therapeutically managing hyperglycemia and associated risk from complications associated with type 2 diabetes mellitus (T2DM). The utility of GLP-1 agonists in promoting weight loss in the setting of obesity has greatly expanded the role of the drug class. Interestingly, over two decades after the FDA approval of exenatide as an adjunctive therapy to improve glycemic control in adults with T2DM, the developmental race to design new drugs and repurpose preexisting therapies targeting the GLP-1 receptor is still underway.

Early studies identified the molecular biology of incretins and the role that endogenous GLP-1 signaling serves in modulating insulin release from beta-islet cells of the pancreas, providing a rational therapeutic basis for the use of these drugs to improve glycemic control [1–6]. However, the much broader GLP-1 receptor expression pattern, including in the central nervous system, suggests that GLP-1 physiology is more elaborate than an endocrine modulator of glycemic control [7–9]. From this perspective, the initially surprising clinical findings that GLP-1 agonists are pharmacologically relevant in targeting disease processes in the heart, vasculature, kidneys, liver, and brain match our developing and inadequate understanding of incretin physiology [10–17]. Further, unexpected findings from retrospective clinical studies suggest that GLP-1 agonists may have additional untapped therapeutic potential in various neurologic and psychiatric conditions [18–20]. However, despite these early signals that GLP-1 agonists may have even broader clinical utility, there remains a significant deficit in our understanding of incretin signaling in the central nervous system. In order to aid drug repurposing efforts attempting to realize the clinical utility of the GLP-1 agonists in the setting of Parkinson’s disease, Alzheimer’s disease, and age-related cognitive decline, a better understanding of the fundamental mechanisms by which GLP-1 agonists affect neural circuits underlying these conditions is required.

Preclinical studies demonstrate that GLP-1 agonists and/or central GLP-1 signaling attenuate neuroinflammation, mitigate tau pathology, improve neurotrophic support, facilitate adult neurogenesis in the hippocampus, and other mechanisms [21–28]. Additionally, multiple lines of evidence in rodent and human studies also suggest that GLP-1 agonists have a highly restricted brain biodistribution pattern [29–32]. Whereas most CNS-active drugs distribute broadly through the brain, certain GLP-1 agonists distribute more selectively into the hypothalamus and specific brainstem nuclei. In most other brain regions, these GLP-1 agonists appear to be below the limit of detection and may not achieve tissue concentrations that are relevant to more broadly engage GLP-1 receptors in the whole brain. As such, reconciling brain wide neuroprotective effects highlighted in preclinical research with the limited brain biodistribution pattern is conceptually problematic. *How can a drug promote effects in a brain region if it does not have access to that region?* This conceptual mismatch between pharmacokinetics and brain wide neuroprotective effects led us to consider that there were unrecognized pathways, cell populations, and even potentially circuits which could facilitate these broad neuroprotective effects remotely.

In this research we identified specific neuronal populations that were differentially activated by semaglutide. Among neuronal populations in the hypothalamus, we considered that orexin neurons of the lateral hypothalamic area exhibit several degrees of functional overlap with the GLP-1 agonists. Further, rodent electrophysiology studies previously identified that bath application of GLP-1 depolarizes and increases the spontaneous firing rate of orexin neurons [33]. Based on this logic, we hypothesized that the orexin system and its neuronal projection targets would be acutely activated by peripherally administered semaglutide. Informed by the results of our histological analysis, we next considered that semaglutide would likely modulate cholinergic neurotransmission in the hippocampus. To test this hypothesis, we employed in vivo microdialysis in young (3-5 months) and aged (23-26 months) rats to identify an acute drug-induced neurochemical effect in the hippocampus. More generally, because this neurochemical effect correlates with learning and memory and is implicated in age-related cognitive decline and neurodegenerative disorders, this research highlights a specific neurochemical mechanism influenced by semaglutide which is relevant to future drug development and repurposing efforts targeting these disorders.

## Results

### Peripherally administered semaglutide coactivates orexin neurons of the lateral hypothalamus and cholinergic neurons in the medial septum of the basal forebrain

To identify specific neuronal populations with altered cellular activity following acute semaglutide treatment, we treated adult rats with either 0.12 mg/kg semaglutide or vehicle (12% DMSO in normal saline) via the intraperitoneal route 2 hours prior to euthanasia. Given that semaglutide concentrates in the hypothalamus and regulates feeding and metabolism, we first assessed if semaglutide would influence the activity of neuronal populations in the hypothalamus that are functionally adjacent to feeding regulation and metabolism.

Within the lateral hypothalamus (LH), we subjectively observed differentially upregulated cFos labeling in orexin neurons of the lateral hypothalamic subdivision (defined as lateral to center of the fornix), but no qualitative difference in the medial bank of orexin neurons (defined as medial to the center of the fornix). Quantitatively, animals that were treated with semaglutide exhibited increased activity in orexin neurons of the lateral hypothalamic subdivision (LHA) regardless of biological sex, as shown in Figure 1A; Main effect of treatment F(1,21) = 13.8, P = 0.0013. Because some clinical effects seen with GLP-1 agonists are dependent on metabolic state of the subject and given that orexin neurons of the LH functionally integrate metabolic information and participate in neuronal circuits that influence feeding and metabolism, we considered the possibility that pre-treatment weight might influence intersubject variability in orexin neuronal activity. However, there was no correlation between pre-treatment weight and orexin neuronal activity (r^2^ = 0.004, P = 0.78), as shown in Figure 1B.

**Figure 1.**
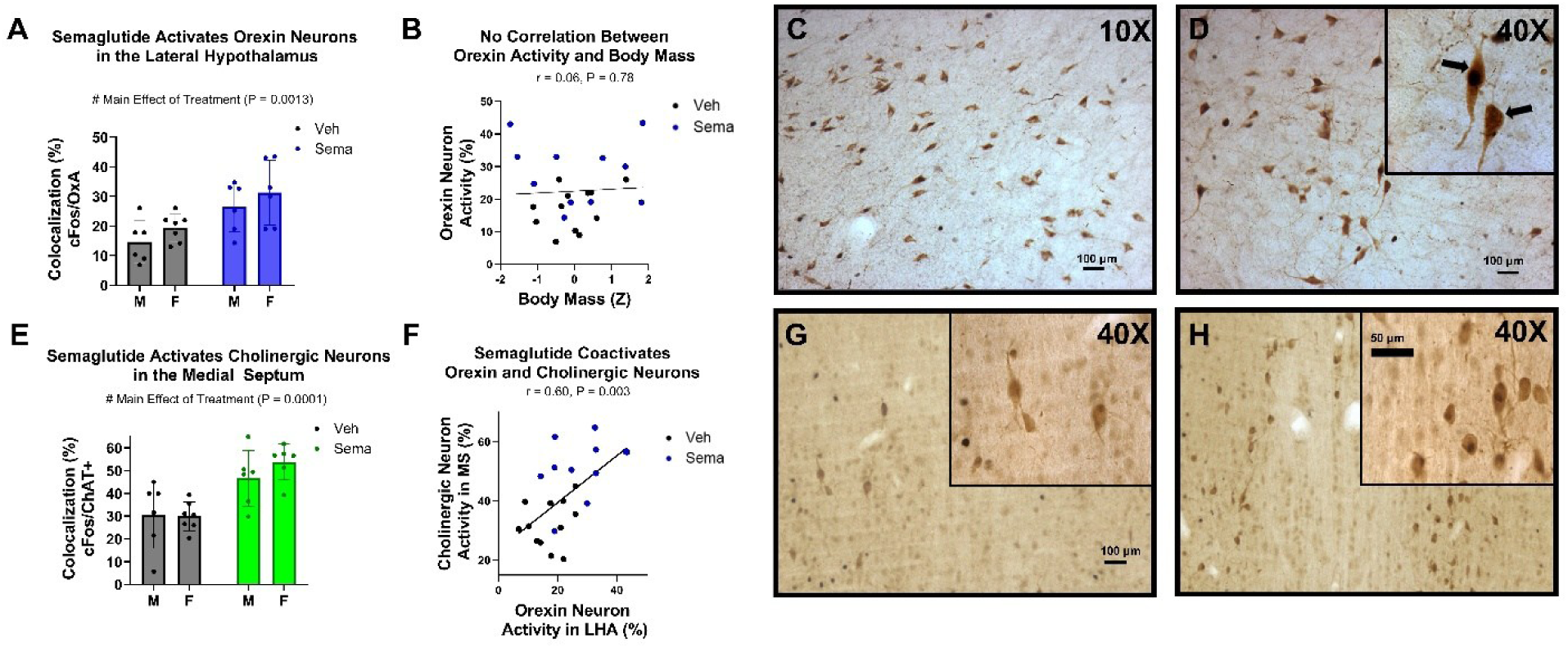
Semaglutide acutely coactivates orexin neurons in the lateral hypothalamic area and cholinergic neurons in the medial septum. Young adult rats received 0.12 mg/kg semaglutide or vehicle (12% DMSO in normal saline) via IP route and were euthanized 2 hours after injection. cFos immunoreactivity within defined neuronal populations was used to assess acute treatment effects on neuronal activity. **(1A)** In the lateral hypothalamic area (defined as lateral to the center of the fornix), semaglutide treatment increased activity of orexin neurons (P = 0.0013) **(1B)** Pre-treatment body weight was not correlated with cellular activity in orexin neurons (P = 0.78). **(1C)** Photomicrographs with orexin labeling (brown) and cFos nuclear labeling (black) in the lateral hypothalamic area in a representative vehicle treated animal. **(1D)** Similarly, a photomicrograph of a representative animal that received semaglutide. The magnified box shows a characteristically double labeled orexin neuron (left arrow) and an orexin neuron without cFos nuclear labeling (right arrow). **(1E)** In the medial septum of the basal forebrain, semaglutide treatment increased activity of cholinergic neurons (P = 0.0001). **(1F)** Across all subjects, activity between population of cholinergic neurons in the medial septum and orexin neurons in the lateral hypothalamic area were correlated (r = 0.c0, P = 0.003). **(1G,H)** Representative photomicrographs of the medial septum with ChAT+ immunoreactivity (brown) and cFos immunoreactivity (black) in vehicle and semaglutide treated animals, respectively. Statistical Analyses: Colocalization data presented were analyzed by 2-Factor ANOVA with treatment and sex as factors. Correlations in B and F were analyzed by simple linear regression (Pearson’s r) with p-values testing if slope is non-zero.

Orexin neurons of the lateral hypothalamus project to a variety of other destinations in the CNS and regulate a broad array of other physiologic processes such as circadian rhythm, social interaction, reward processing, and others [34]. However, orexinergic projections to the basal forebrain are especially functionally relevant to cognition and both orexin and basal forebrain cholinergic systems are compromised in aging and Alzheimer’s disease [35–40]. Because of structural relevance and translational significance, we considered that increased activity in the orexin system may coincide with increased activity in the basal forebrain cholinergic system (BFCS)—even if semaglutide presumably does not distribute to this brain region. Subjectively, we observed a qualitative increase in cholinergic neuron activity in animals that received semaglutide in only the medial septum subregion of the BFCS. Quantitatively, animals that were treated with semaglutide exhibited increased activity in cholinergic neurons of the medial septum subregion regardless of biological sex, as shown in Figure 1E; F(1,21) = 21.9, P = 0.0001. Additionally, we found that activity between orexin neurons of the lateral subdivision of the LH and cholinergic neurons within the medial septum subdivision of the basal forebrain was globally correlated across all subjects, as shown in Figure 1F (r^2^ = 0.36, P = 0.0026). Taken together, these histological results suggest that acute treatment of semaglutide coactivates two structurally and functionally interconnected neuronal populations in young adult rats.

### Peripherally administered semaglutide acutely increases acetylcholine efflux in the ventral hippocampus

The projection field of the BFCS is determined by subregion and the medial septum predominately sends its projections broadly throughout the hippocampus via the septo-hippocampal pathway. Given that medial septum cholinergic neurons were acutely activated by semaglutide and when considering age-related changes that occur in both the orexin and basal forebrain cholinergic systems, we next hypothesized that semaglutide may increase cholinergic transmission in the hippocampus of young but not aged animals.

To test this hypothesis, we employed in vivo microdialysis to assess acetylcholine efflux in the ventral hippocampus of young (3-5 months) and aged (23-26 months) rats treated with both semaglutide and vehicle in a crossover design study with a 48-hour washout period between treatments. The 48-hour washout period was chosen because of the shortened pharmacokinetic elimination half-life of semaglutide in rats versus humans. In our cross-over microdialysis study, we assessed intersession weight loss over 48 hours and observed a main effect of treatment, as shown in Figure 2C; Effect of treatment, F(1,36) = 44.2, P < 0.0001; Sex x Age Interaction, F(1,36) = 8.3, P = 0.0068; Sex x Age x Treatment Interaction, F(1,36) = 11.3, P = 0.0019. These confirmatory results suggest that the dose, route, and carrier formulation used with semaglutide in this study were producing known physiologic effects associated with GLP-1 agonists in young and aged rats. Cross-over studies permit comparison of treatment effects within subjects if sufficient washout occurs between treatments. However, the assumption that the 48-hour washout between each session was sufficient for a cross-over study was proven to be false. We assessed baseline acetylcholine levels across microdialysis sessions and found a significant treatment-session interaction in young animals and a sex-treatment-session interaction within aged animals, as shown in Supplemental Figure 1. In young adult animals, treatment with semaglutide in the first session resulted in increased baseline levels in the second session; Effect of treatment, F(1,15) = 10.3, P = 0.0059; Treatment x session interaction, F(1,15) = 10.3, P = 0.0059. Unlike in young animals, these differential carryover effects across microdialysis sessions were more complex in aged animals as they were also influenced by sex; Sex x treatment interaction, F(1,16) = 9.4, P = 0.0075; Sex x treatment x session interaction, F(1,16) = 9.4, P = 0.0075. To overcome the differential carryover effect within our cross-over study design and avoid erroneously using false-control groups in the second session, we analyzed acetylcholine efflux using a between groups analysis within the first session only. We chose to analyze the first session only because of the concern in the difficulty of interpreting the second session in isolation due to the concern of lingering drug effects across sessions.

**Figure 2.**
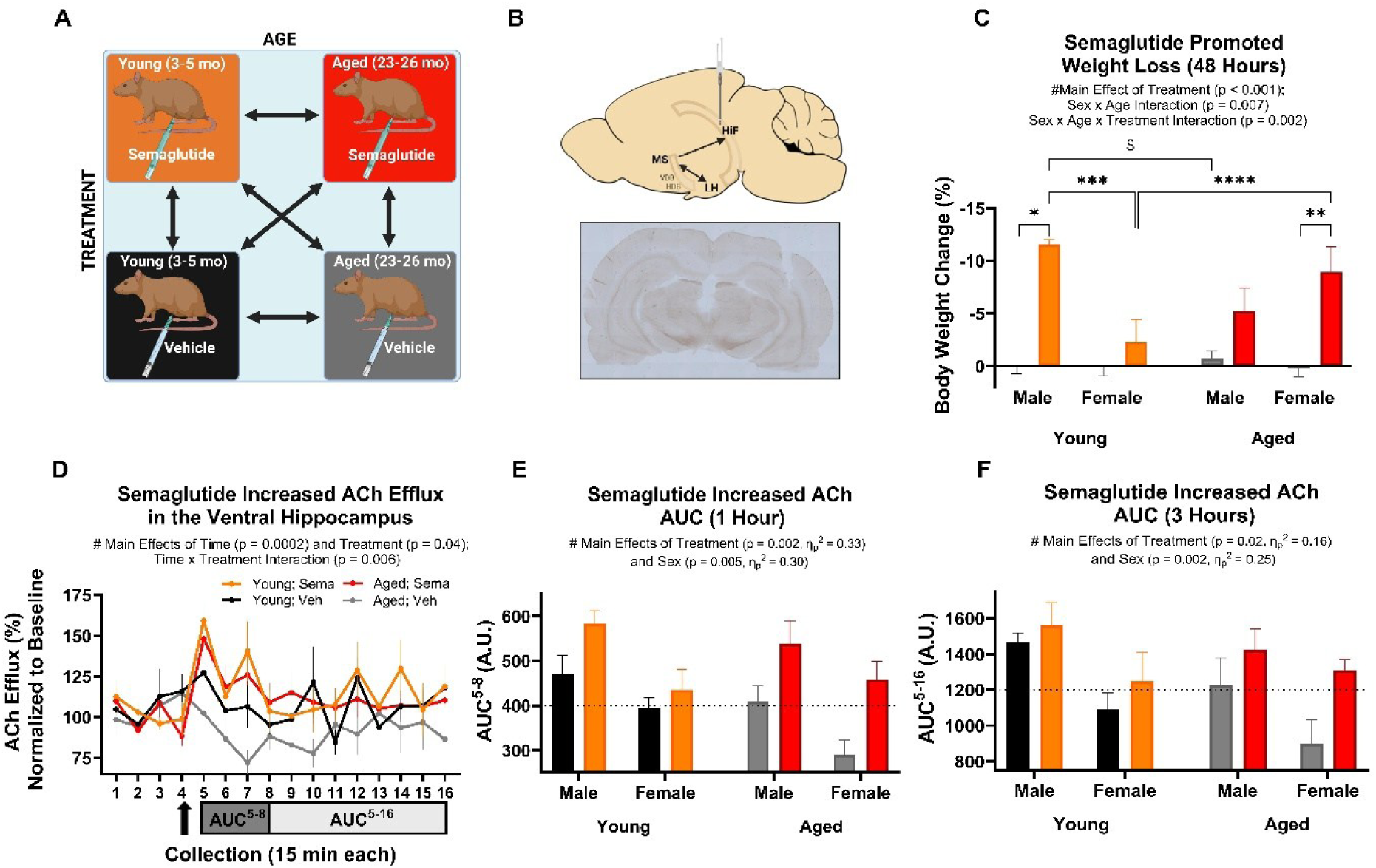
Semaglutide acutely increases acetylcholine efflux in the ventral hippocampus of young and aged rats. In vivo microdialysis was performed in freely moving young adult (3-5 mo) and aged (23-2c mo) rats to assess acetylcholine efflux in the ventral hippocampus in response to treatment with semaglutide or vehicle. **(A)** Within the first session, young adult or aged rats, balanced for sex, received either 0.12 mg/kg of semaglutide or vehicle (12% DMSO in normal saline) via IP route. **(B)** Conceptualization of circuit-based hypothesis that semaglutide would increase in acetylcholine efflux in the ventral hippocampus and representative tissue with canula and probe target location. **(C)** As a confirmatory measure, weight loss was tracked over 48 hours and was inffuenced by treatment (P < 0.0001). In addition, there was a sex x age interaction (P = 0.007) and a sex x age x treatment interaction (P = 0.0002). Statistically significant pairwise comparisons between groups are indicated (adjusted P < 0.05). **(D)** Graphical representation of acetylcholine (ACh) efflux over time within the first session where each collection represents a 15 min time interval. ACh concentration in dialysate collections were normalized to their baseline, which was calculated as the median of the first 4 basal collections. Graphical time-series data represents biological sex as a pooled variable. Animals received either semaglutide or vehicle after the last basal collection as indicated by the arrow. In this analysis across all collections, there were statistically significant effects of time (P = 0.0002), treatment (P = 0.04), and a time x treatment interaction (P = 0.008). **(E, F)** Additionally, two AUC analyses representing cumulative normalized ACh efflux over 1 hour and 3 hours following treatment are shown. In the AUC (1 hour) analysis, there were significant effects of treatment (P = 0.002) and sex (P = 0.005). In the AUC (3 hours) analysis, there were significant effects of treatment (P = 0.02) and sex (P = 0.002). Statistical Analysis: ACh efflux time series data shown as mean +/− SEM for each group and statistical analysis was performed using mixed effects analysis with time, treatment, and age as variables. In both AUC analysis, group data is depicted as mean +/− SEM and statistical analysis was performed by two separate 3-Factor ANOVAs with treatment, age, and sex as variables.

Within the first microdialysis session only, normalized levels of acetylcholine efflux in the ventral hippocampus were collected in 16 aliquots acquired over 4 hours, with animal subjects receiving either semaglutide or vehicle immediately after collection 4, as shown in Figure 2D. In the first primary neurochemical outcome of the study, we assessed acetylcholine efflux in the ventral hippocampus by a mixed effects model with treatment, time, and age as variables of interest with experimental groups pooled and balanced for sex. In this analysis, there were main effects of treatment, time, and a time x treatment interaction; Treatment F(1, 38) = 4.37, P = 0.043; Time F(15, 567) = 2.86, P = 0.0002; Time x Treatment Interaction F(15, 567) = 2.18, P = 0.0062. For the second primary neurochemical outcome of the study, we performed two area under the curve (AUC) analyses at time intervals of biological significance within the first microdialysis session. These AUC analyses represent cumulative acetylcholine efflux over 1- and 3-hour time intervals after treatment and were analyzed by 3-Factor ANOVA with treatment, sex, and age as variables of interest. Across all groups pooled for age and sex, semaglutide treatment resulted in a 27.7% relative increase in the 1-hour AUC for acetylcholine as shown in Figure 2E. In the AUC (1 hour) statistical analysis there was a main effect of treatment, F(1,34) = 16.8, P = 0.0002, and sex, F(1,34) = 14.9, P = 0.0005. In the AUC (3 hour) analysis shown in Figure 2F, there was also a main effect of treatment, F(1,34) = 6.5, P = 0.016, and sex, F(1,34) = 11.2, P = 0.002. Together, these results indicate that semaglutide treatment acutely increased acetylcholine efflux in the ventral hippocampus, regardless of age or sex.

## Discussion

In this study, we identified orexin neurons of the lateral hypothalamus as a specific neuronal population that exhibited increased activity following acute semaglutide administration in young adult rats. To our knowledge this is the first study to report this finding and one of the first studies to identify a specific neuronal population whose activity is influenced by any GLP-1 agonist. In support of these results, two rodent biodistribution studies using whole brain clearing and light sheet microscopy or tissue mass spectrometry found that semaglutide concentrates specifically in the hypothalamus [29, 31]. In support of the framework that GLP-1 agonists may drive central effects directly through distribution and receptor engagement in the hypothalamus, GLP-1R mRNA is abundantly expressed in the hypothalamus and rodent electrophysiology studies using acute slice preparations show that bath application of GLP-1 depolarizes and increases the spontaneous firing rate or orexin neurons in the LH [33]. However, our study does not specifically demonstrate that semaglutide directly engages GLP-1 receptors expressed by orexin neurons. Rather, Fos immunoreactivity is a general measure of cellular activity and cFos is expressed downstream from GPCR signaling and other signaling pathways. Importantly, a further consideration when interpreting this result is that other neuronal populations and/or metabolic information influences orexin activity in the context of hypothalamic circuitry. For example, orexin neurons directly sense glucose availability via intracellular pools of purinergic nucleotide metabolites and responsively depolarize and increase their spontaneous firing rate in hypoglycemic conditions [41, 42]. Therefore, an alternative rationale that might explain our results that semaglutide increased orexin neuron activity in the lateral hypothalamus is that the hypoglycemic effects of semaglutide could depolarize and generally activate orexin neurons through this glucose sensing mechanism characteristic of orexin neurons. In addition, because there is also evidence of GLP-1 agonist biodistribution into specific circumventricular organs of the brainstem, it is reasonable to assume that direct effects in the brainstem might indirectly mediate increase orexin activity in the hypothalamus through a circuit mechanism. For these reasons and because the orexin system is an interesting pharmacologic target in several neurological disorders, it would be translationally meaningful to investigate exactly how orexin neurons are activated by peripherally administered semaglutide. Here, electrophysiology studies using acute slice preparations would be inherently confounded by bypassing the unique constraints imposed by the limited biodistribution of GLP-1 agonists entirely. An elegant alternative approach would be to use fiber photometry with specific probes that report GLP-1R engagement (such as GLPLight1) in vivo [43]. This experimental approach would allow for monitoring GLP-1R activation on orexin neurons in behaving rodents without significantly disrupting the blood-brain barrier. However, to our knowledge, while GLPLight1 has been validated in vitro, this probe has not been validated in rodents at this time.

While it was expected that increased orexin neuron activity would also influence cholinergic activity in the basal forebrain due to their structural and functional connectivity, there are three further considerations worth noting. First, there is no current evidence at the time of this study that suggests that semaglutide achieves pharmacologically meaningful concentrations in the medial septum that could directly activate cholinergic neurons. Secondly, referencing the Allen Brain Atlas for murine in situ hybridization data for GLP-1R expression shows that GLP1R mRNA expression is highly abundant in the dorsolateral septum—but sparse or even absent in the medial septum [44]. This implies that the increased activity seen in cholinergic neurons of the medial septum is likely an indirect effect which is not directly facilitated by GLP-1 receptor engagement and subsequent cell signaling in cholinergic neurons residing within the medial septum. A limitation to this logic is that spatial patterns of GLP-1R expression are not necessarily conserved across rodent species. The last consideration regarding this result is that it is not clear why we saw qualitatively increased activity in the medial septum specifically as opposed to a global increased activity within cholinergic neurons across all basal forebrain subregions. One possible explanation for this observation could be that only specific subpopulations of orexin neurons are activated following acute semaglutide treatment. In support of this notion, it has been shown via laser-capture microdissection that orexin neurons of the LH in rodents are a genetically heterogenous cellular population [45]. To our knowledge, it has not been shown if any specific clusters or subpopulations of orexin neurons specifically express GLP-1R or exhibit a bias in their projections to the medial septum. However, one possible explanation for the spatially selective pattern of activation in the basal forebrain is that semaglutide may activate specific orexin neurons which more selectively project to medial septum neurons.

The findings that semaglutide increased activity in the orexin and basal forebrain cholinergic systems alongside increased cholinergic efflux in the hippocampus are fascinating given that each of these systems is dysfunctional in aging and dementia. In humans, orexin neurons of the lateral hypothalamus are reduced and/or phenotypically silenced in aging and Alzheimer’s disease. In parallel to these findings, our research group has previously shown an age-related decline in orexin immunoreactivity in the strain of rats used in this work [40]. Interestingly, the magnitude of the reduction seen in orexin-A immunoreactive cells in these Brown Norway F1 x F344 hybrid rats is quantitatively similar to the reduction seen in post-mortem studies of aged human subjects [39]. However, it remains uncertain whether this loss of immunoreactivity to orexin neuropeptides is explained by neurodegeneration or if orexin expression is simply silenced, or both in these contexts.

In the process of aging, the basal forebrain cholinergic system also exhibits functional changes. Specifically, BFCS neurons exhibit altered electrophysiologic properties such as enhanced excitability, presumably driven by increased susceptibility to oxidative stress and neuroinflammation that occur in aging [46]. Taken together, our histologic data demonstrates that semaglutide modulates the activity of neuronal populations that are vulnerable to aging and neurodegeneration and that these systems are important to consider in the context of future preclinical CNS research with any GLP-1 agonist.

Despite the extensive clinical utilization of GLP-1 agonists for over two decades, the finding that peripherally administered semaglutide produces an acute increase in acetylcholine efflux within the ventral hippocampus is among the first specific neurochemical effects demonstrated with the GLP-1 agonists. Prior to this study, it has also been shown that dopamine efflux in the nucleus accumbens is blunted when semaglutide is given as a pre-treatment before cocaine [47]. While blunted dopamine release provides powerful translational insight into the mechanism through which GLP-1 agonists produce clinical effects in substance use disorders and perhaps hedonic feeding regulation, enhanced acetylcholine efflux in the hippocampus provides specific neurochemical insight into how GLP-1 agonists may function in the context of age-related cognitive decline, Alzheimer’s disease, and related dementias.

Deeper insight into the implications of this neurochemical effect is found when considering the established physiologic role of cholinergic signaling in the hippocampus. In the hippocampal formation, cholinergic neurotransmission supports long-term potentiation, supports theta-rhythmicity, suppresses neuroinflammation, and may support adult neurogenesis [48–51]. All these effects have also been demonstrated in preclinical models with GLP-1 agonists with the caveat that GLP-1 agonists do not appear to distribute into the hippocampus. Therefore, it is likely that GLP-1 agonists indirectly mediate these hippocampal effects through other systems. Given our study findings that semaglutide acutely increased acetylcholine efflux in the hippocampus, it is possible these established hippocampal effects associated with the with the GLP-1 agonist therapeutic class are indirectly facilitated via cholinergic signaling.

Congruently, deterioration of the cholinergic system produces impairment in learning and memory. As a particularly relevant example, local injection of an orexin-saporin toxin into the medial septum and diagonal band of Broca disrupted spatial reference memory and spatial working memory in rats [52]. This particular experimental model produced degeneration of cholinergic and GABAergic neurons in the medial septum and diagonal band of Broca that also express orexin receptor type-II. Selective degeneration of these orexin receptor expressing neurons in this model provides behavioral insight into orexin-basal forebrain-hippocampal circuitry and provides perspective on the results of this study. That is, given that semaglutide concentrates in the hypothalamus and based on our results that semaglutide coactivated orexin neurons in the lateral hypothalamus and cholinergic neurons in the medial septum and acutely increased acetylcholine efflux in the hippocampus, it is possible that semaglutide modulates a hypothalamic-basal forebrain-hippocampal circuit that supports learning and memory. However, while the results of this study suggest this circuit mechanism, this study was not designed to definitively prove a circuit mechanism. Rather, the intent of this microdialysis experiment was to evaluate if semaglutide would increase cholinergic efflux in the hippocampus and how age and biological sex would influence this drug-induced neurochemical effect.

Our hypothesis was that age would determine the acute neurochemical response to semaglutide. We considered that aged animals would exhibit a blunted neurochemical effect because of age-related changes in the orexin and basal forebrain cholinergic systems. We also considered that biological sex could influence the neurochemical effect given that orexin neuropeptide expression is regulated by estrogen signaling [53, 54]. After we designed and completed this microdialysis study, a translational neuroscience study was published which also suggests that metabolic dysfunction and biological sex determine GLP-1 agonist uptake in the human brain. Specifically, a radiolabeled GLP-1 agonist achieved higher uptake (SUV_max_), in the pituitary of male patients with diabetes [32]. Although there are multiple theoretical mechanisms through which age and/or biological sex could influence drug effects, our study shows that semaglutide globally increased acetylcholine in the hippocampus at 1 and 3 hours after treatment regardless of age and sex. The finding that age did not acutely diminish semaglutide-induced acetylcholine efflux in the hippocampus is translationally relevant when considering the patient population that would receive GLP-1 agonists in the context of dementia or cognitive decline.

More broadly, semaglutide appears to have functional overlap with the cholinergic hypothesis of dementia. Arguably, while no single mechanism can independently account for Alzheimer’s disease and related dementias, the cholinergic system is intimately connected with other relevant systems and pathomechanisms that are implicated in the development and progression of dementias. Accordingly, pharmacologically inhibiting acetylcholinesterase to limit synaptic metabolism of acetylcholine and extend cholinergic signaling from a deteriorating cholinergic system has been the primary therapeutic strategy employed in managing dementia for several decades. However, this measure has been insufficient and does not dramatically alter disease trajectory but primarily serves to manage symptoms. Here, an important general consideration is that drug mechanisms that increase neuronal activity and neurotransmitter release are distinct but also sometimes synergistic with drug mechanisms which inhibit enzymatic synaptic transmitter degradation. This present study found that semaglutide acutely facilitated a modest increase in acetylcholine efflux in the ventral hippocampus. However, the possibility of pharmacodynamic synergy between GLP-1 agonists and systemically administered acetylcholinesterase inhibitors is a highly translationally relevant future research direction given the prevalence of acetylcholinesterase inhibitor use and when considering that many dementia clinical trials provide experimental therapeutics in addition to standards of care.

Besides “probing” acetylcholinesterase inhibitor-GLP-1 agonist pharmacodynamic interactions, there are additional pathophysiologic variables that may provide deeper translational insight in future research directions related to this study. First, it may be that neuroinflammation is an important variable that modifies the significance of semaglutide-mediated increases in cholinergic neurotransmission. GLP-1 agonists and hippocampal cholinergic transmission both produce anti-inflammatory effects. In contrast, AD-related molecular pathology produces hippocampal inflammation that drives cognitive dysfunction in various animal models. Therefore, AD molecular pathology and the resulting hippocampal neuroinflammation that it produces may specifically highlight the therapeutic potential of GLP-1 agonists. Secondly, our study did not find any meaningful correlation between pretreatment weight and any effect. However, it is likely that metabolic dysfunction is an important variable underlying neuroprotective effects of the GLP-1 agonists. In late 2025, the evoke and evoke+ clinical trials were discontinued because they did not find a clinically significant neurocognitive benefit with oral semaglutide treatment in a mild-cognitive impairment (MCI) and mild-AD patient population that was mostly non-diabetic [55]. In contrast, a large meta-analysis of prospective trials published the same year found a protective effect in a general diabetic patient population that were treated with GLP-1 agonists. Importantly, this same meta-analysis found no benefit in protecting against dementia or cognitive decline with the sodium-glucose cotransporter (SGLT-2) inhibitor therapeutic class [56]. These clinical data suggest that (1) there may be a therapeutic class-specific effect among type II diabetes mellites patients and (2) that there may be critical metabolic determinants of clinical effectiveness when repurposing the GLP-1 agonists for dementia and cognitive decline. As such, a critical preclinical research question that remains is whether there are certain metabolic variables that might unmask therapeutic susceptibility to cognitive benefits with the GLP-1 agonists in AD models and in the context of age-related cognitive decline.

This study sought to identify specific neuronal populations and neurochemical mechanisms that could aid in repurposing the GLP-1 agonist therapeutic class for CNS disorders. The findings that semaglutide increased activity in the orexin and basal forebrain cholinergic systems alongside increased cholinergic efflux in the hippocampus provide fundamental insight into how the therapeutic class may target specific functional deficits that occur in the setting of aging and neurodegenerative disorders.

## Methods

### Animals

All experiments were conducted according to protocols drafted with guidance from the National Institutes of Health Guide for the Care and Use of Laboratory Animals and approved by the University of South Carolina Institutional Animal Care and Use Committee (protocol 2706-101 900-022824). Fisher 344/Brown Norway F1 hybrid rats, from the National Institute on Aging colony (Baltimore, MD, USA) were used for both histology (age 3-5 months) and microdialysis experiments (ages 3-5 months and 23-26 months). All experimental groups included equal numbers of male and female rats. Animals were housed with *ad libitum* access to food and water in a climate-controlled animal facility operating on a 12/12 hr light cycle. All experiments were conducted during the light phase.

### Acute Semaglutide Histology Experiment

For acute histology experiments, animals were habituated to transport between home and study environments, study room, and handling for 3 days prior to the experimental day. On experimental days, animals were brought to an acoustically isolated room and allowed to acclimate for 30 min. Then, animal subjects were provided with either 0.12 mg/kg semaglutide (AA BLOCKS, INC. Lot # AAB25-643507-1) or an equivalent weight-based volume of vehicle (12% DMSO in normal saline) administered intraperitoneally. Rats exhibit faster clearance of semaglutide than humans and this dose was selected because it represents a behaviorally and physiologically relevant dose as surmised from the rodent literature [57–59]. After receiving their injection, animals were returned to their cages with *ad libitum* access to food and water. Experimental days were broken up into animal cohorts of up to 8 animals that were balanced for sex and treatment. On experimental days, animals received semaglutide or vehicle on a sequential (rolling) basis at 30 min intervals starting at 2-3 ZT and ending by 5-6 ZT. Exactly 2 hours after injection, animals were anesthetized under inhaled isoflurane immediately prior to transcardiac perfusion with 200 mL of phosphate buffered saline and 300 mL of 4% paraformaldehyde. Whole brains were removed and stored in 4 % paraformaldehyde at 4 C until further histology processing.

### Histology and Microscopy

Brains were cut into 50-micron thick coronal sections with a vibratome and immunohistochemistry in free floating sections was performed and included the following preincubation steps: washing in TBS buffer, endogenous peroxidase quenching with dilute hydrogen peroxide (3% in TBS), and blocking/permeabilization in horse serum with dilute triton-X. Double-labeled IHC was performed using primary antibodies directed against rat cFos and either mouse anti-orexin-A or goat anti-choline acetyltransferase (ChAT), followed by biotinylated or unlabeled secondary antibodies and streptavidin horseradish peroxidase or species-appropriate peroxidase anti-peroxidase, respectively. Tissue sections were then stained using 3,3’-diaminobendizine (DAB) and hydrogen peroxide with development times that were subjectively optimized under a light microscope to achieve optimal labeling for each label. Nickel-cobalt-enhanced DAB was used to visualize black/blue cFos-positive nuclear staining. Following c-Fos labeling, neuronal identity labeling was performed and developed in plain DAB in order to identify cholinergic (ChAT-positive) or orexinergic (OxA-positive) brown cell bodies. After mounting, dehydration/delipidation and cover slipping, cell counting was performed manually on a Nikon Eclipse E600 light microscope at 20-40X magnification. Photomicrographs were acquired on a Nikon Eclipse E600 or an ECHO Revolution microscope.

Preclinical research identifying which neuronal populations are activated or inactivated by GLP-1 agonists is limited. Because drugs can influential neuronal activity in a spatially selective manner, counting all neurons in a brain region could dilute meaningful drug effects. To overcome this concern, we used a qualitative approach to refine our hypothesis to specific neuroanatomical subregions and subsequently quantified neuronal activity in specific neuronal populations within specific subregions as defined by Paxinos and Watson.

### Antibodies

Rabbit anti-cFos (Millipore-Sigma ABE 457, Lot 3585299) was used at a 1:5000 dilution. Mouse anti-orexin A was used at a 1:500 dilution (Santa Cruz KK09, Lot SC-80263). Goat anti-choline acetyltransferase (Millipore-Sigma AB 144, Lot 4020102) was used at a 1:1000 dilution. All secondary antibodies (e.g., biotinylated donkey-anti-rabbit, unlabeled donkey-anti-goat, and unlabeled donkey-anti-rabbit) were obtained from Jackson Immunoresearch. All peroxidase conjugates such as streptavidin-conjugated horseradish peroxidase (SHRP), goat peroxidase-anti-peroxidase, and mouse peroxidase-anti-peroxidase were obtained from Jackson Immunoresearch.

### Histological Measures and Statistical Analyses

Colocalization was determined by first counting all OxA- or ChAT-positive neurons within a defined region of the hypothalamus or basal forebrain, respectively, and then counting how many of those cells had a Fos-positive nucleus. Colocalization was defined by the presence of a blue/black nucleus and contiguously surrounding brown cytoplasm, all within the same plane of focus. Cells were counted across 2-3 representative tissue sections for each animal. Representative tissue sections were randomly selected from a pool of eligible sections and eligibility was determined by the following criteria: appropriate neuroanatomical location, sufficient tissue quality and labeling quality to support cell counting, and sufficient cellular abundance to support robust quantification. Colocalization data is displayed as a bar graph with each individual value representing the mean percent of defined neurons that colocalized with cFos nuclear staining for an individual animal. These data were analyzed by way of 2-Factor ANOVA with sex and treatment as factors with two levels each. Coactivation data and baseline weight vs. orexin activity correlation data is displayed as a scatter plot and simple linear regression was performed.

### Acute Semaglutide In Vivo Microdialysis Cross-Over Experiment

For microdialysis experiments, all animals underwent a sequence of canula placement surgery (day 1), recovery (day 2), habituation (days 3-6), and then two sessions of microdialysis separated by a rest day (days 7-9). These experiments were run in sequential cohorts of 2-6 animals, counterbalanced by treatment sequence and sex. On surgery days, animals received induction anesthesia with 4% isoflurane in an induction box, and then maintenance anesthesia at a reduced flow rate with 4% isoflurane in a stereotaxic apparatus. Microdialysis cannulas (BASi, Inc., West Lafayette, IN) were placed in the ventral hippocampus, counterbalanced by hemisphere, at surgical coordinates (A/P - 5.2 mm, L +/− 5.0 mm, D/V – 4.0 mm) relative to Bregma. Topical anesthesia (intradermal carbocaine/lidocaine), fluid replacement (3 mL subcutaneous normal saline), heat therapy, and post-surgical analgesia (0.05 mg/kg subcutaneous buprenorphine) was provided to all animals. After an additional day of recovery after surgery, animals were brought to a small, visually and acoustically isolated microdialysis room with multiple large microdialysis bowls (BASi, Raturn®). Our microdialysis setup visually separates rodents from each other and permits free rotational movement and allocentric spatial exploration/navigation by sensing relative head position and responsively actuating bowl rotation. Animals received habituation to handling, bowls, and tethering for 4 days and injection (normal saline) for 3 days prior to the first microdialysis session. A cross-over design was used on experimental days such that each animal received semaglutide in one session and vehicle in the other session, counterbalanced for treatment sequence. Microdialysis experiment days occurred exclusively in the light-phase and began at ZT 1-2 and finished by ZT 9-10. During this time interval, animals were restricted to food and water. Microdialysis probes (BASi) were perfused with artificial cerebrospinal fluid at a flow rate of 2 uL/min. After probes were in vitro validated for flow rate, cannula stylets were removed and replaced with probes at ZT 2-3. After in vivo verification of proper flow rate, discard dialysate was collected from each animal for 3 hours. Following the discard period and starting at ZT 5-6, 16 experimental collections (15 min per collection) were acquired. Four baseline dialysates were collected followed by treatment with semaglutide (0.12 mg/kg via intraperitoneal route) or vehicle (12% DMSO in normal saline via intraperitoneal route). Collection resumed immediately following treatment and continued for 3 additional hours (collections 5-16). Dialysates were immediately frozen on dry ice as they were acquired and then stored at −80 C until neurochemical analysis was performed. To control for sample degradation, all dialysates were thawed for 20 minutes prior to neurochemical analysis using HPLC and no sample was refrozen.

### Neurochemical Measures and Statistical Analyses

An HTEC-510 HPLC (Amuza) with an AC-Gel reverse phase column (Eicom), a post-column immobilized enzyme reactor, AC-ENZYM II (Eicom) and an electrochemical detector consisting of a platinum working electrode against an AgCl reference electrode was used to quantitate acetylcholine efflux from microdialysate collections as described previously [60]. A potassium phosphate buffered mobile phase with EDTA and sodium dodecanesulfonate was used for HPLC. The HPLC conditions used for the analysis involved a flow rate of 150 uL/min and an applied potential of 400 mV against a AgCl reference electrode. Every day of neurochemical analysis, a 3-point, 2-analyte standard curve was constructed for choline and acetylcholine from serially diluted stock standard solutions stored at −80 C. In order to mitigate and control for sample degradation, each sample was thawed for 20 min and mixed prior to manual injections. Additionally, the sequence of collections analyzed for a particular animal session was randomized on each day to reduce the risk of introducing bias.

Neurochemical analyses include (I) analysis of basal ACh concentrations between both sessions, (II) baseline ACh levels from session 2 that were normalized to session 1 median basal concentration, (III) ACh levels of each collection in session 1 normalized to session 1 median basal concentration, and (IV) area under the curve (AUC) analyses for ACh in session 1 which consists of a sum of normalized ACh levels over a defined bin of sequential collections (e.g. AUC^5-8^ represents area under the curve for 4 collections over 1 hour immediately after treatment). (I) Basal ACh concentrations are shown as bar graph with each individual data point representing the median of 4 basal concentration measurements for an animal error bars representing standard error of the mean (SEM). (II) Baseline ACh levels normalized to session 1 are shown as a bar graph with each individual data point representing the median of 4 basal concentrations measurements for a given session normalized to the basal ACh concentration from the first session for a particular animal. Error bars represent SEM. A 3-Factor ANOVA was performed with biological sex, treatment, and session as factors; each with 2 levels. (III) ACh levels of each collection in session 1 normalized to session 1 median basal concentration are displayed as time series data for each group with each data point representing group mean and error bars representing SEM. A 3-Factor ANOVA was performed in groups with pooled sex, and factors of time, treatment, and age. Groups were displayed and statistically analyzed with groups that were pooled for sex to improve interpretability and based on the *a priori* assumption that an age-treatment and not a sex-treatment interaction would occur. (IV) Subsequently, both AUC analyses were performed as separate analyses targeting pharmacologically relevant time epochs that capture an acute drug effect. AUC analyses are displayed as bar graphs with each data point representing the AUC over a defined time interval for a particular subject. In these AUC analyses, time was collapsed into a single level—which permitted use of a 3-Factor ANOVA with treatment, age, and sex as variables. Because two AUC time intervals were chosen and two separate ANOVAs were run, we did not assume that a significant omnibus ANOVA F-test was sufficient to support group differences over both AUC analyses. Instead, all AUC effects and interactions from the two related ANOVAs were combined into a single predefined family, and P-values were adjusted using the Benjamini–Hochberg step-up procedure to control the false discovery rate at q = 0.05.

### Outliers and Excluded Data

No outliers were identified and no data was excluded for histology experiments. In neurochemistry experiments, 2 animals were excluded due to probe placement being outside of the target area, and 4 animals in unique groups were identified as outliers and excluded from specific statistical analyses (as discussed below). <u>Data excluded from neurochemical analyses due to experimental reasons:</u> In order to permit meaningful comparisons across sessions, all data from both sessions were excluded from analysis for a particular animal if dialysis probe failed as evidenced by stopped flow or reduced output during experimental collections from either session. In addition, all data was excluded from a particular animal if probe placement was outside of the target area and all data from two animals were excluded for this reason. <u>Data excluded from neurochemical analyses due to statistical reasons:</u> The neurochemical analyses (I) and (II) had 4 animals from different groups considered as outliers and their data was excluded using conventional statistical criteria (1.5IQR < Q1 or > Q3). No data was excluded from the primary neurochemical analyses (III) and (IV). Outliers were excluded from analyses (I) and (II) using statistical criteria because these analyses were only used to determine how the primary neurochemical analyses were analyzed. The implication of excluding these 4 animals is that it led us to abandon a within-subjects analysis in favor of a between groups analysis in the primary neurochemical analysis. While this more conservative approach reduced statistical power in the primary neurochemical analysis, it also likely helped us detect and avoid erroneously using a within-subjects design that would have been confounded by unexpected prolonged drug effects and an inadequate washout period between microdialysis sessions.

## Author Contributions

CW designed research studies, conducted experiments, acquired data, analyzed data, and wrote the manuscript. JF designed research studies, conducted experiments, and wrote and edited the manuscript. AB conducted experiments, acquired data, and analyzed data. MF and JW conducted experiments and edited the manuscript. NM and AP acquired data. CG and LR designed research studies and edited the manuscript.

## Acknowledgements

We would like to thank the University of South Carolina School of Medicine Instrument Resource Facility for training and support with microscopy. Special thanks to Matthew Efird and Mike Gore of the Pharmacology, Physiology and Neuroscience Machine Shop for technical assistance with research equipment and instrumentation.

Financial support for these studies was provided by the National Institutes of Health (RO1/2RF1 AG050518 to JF), and by the Office of the Vice President for Research at the University of South Carolina (SPARC 180800-25-705022 to CW)

