## Supplemental Figure 1 for "Semaglutide Activates the Orexin/Hypocretin and Basal Forebrain Cholinergic Systems and Increases Acetylcholine Levels in the Hippocampus of Young and Aged Rats"

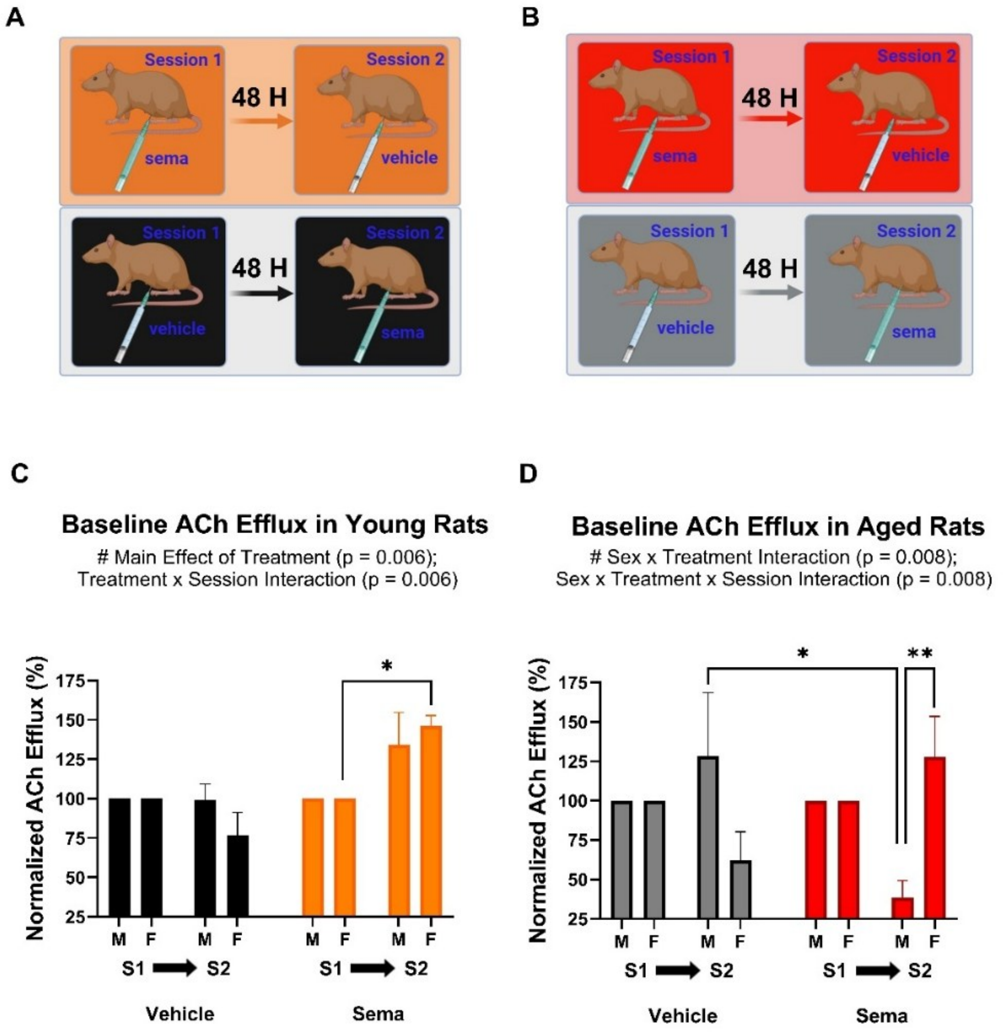


**Supplemental Figure 1: Crossover design and baseline session characteristics of in vivo microdialysis experiment.** ***(A, B)*** *Crossover design involved two sessions separated by 48 hours of young (A; 3-5 months) or aged (B; 23-26 months) rats. Groups were balanced for sex and treatment sequence. We analyzed session baseline levels within subjects to determine if session could appropriately be treated as a within-subjects variable for the primary neurochemical analysis.* ***(C)*** *In young animals, there was a main effect of treatment (P = 0.006) and a treatment x session interaction (P = 0.006).* ***(D)*** *In aged animals, there were statistically significant interactions including sex x treatment (P = 0.008) and sex x treatment x session interaction (P = 0.008). These results indicate the presence of a differential carryover effect across sessions in both young and aged groups and were used to guide analysis of subsequent primary neurochemical measures. Due to the differential carryover effect, we concluded that analyzing the primary neurochemical outcome data with session as a within subjects variable would be inappropriate since semaglutide given in the first session appeared to have altered acetylcholine efflux in the second session despite treatment with vehicle. Statistical Analyses: Baseline acetylcholine level data across sessions for a specific animal were calculated as the median of 4 basal levels in each session normalized to the session 1 baseline for each animal. Group data is represented as mean +/- SEM and was analyzed by 3-Factor ANOVA with treatment, sex and session as factors.* All pairwise comparisons (adjusted P < 0.05)
